# The secretome of M1 and M2 microglia differently regulate proliferation, differentiation and survival of adult neural stem/progenitor cell

**DOI:** 10.1101/2021.09.15.460424

**Authors:** Xue Jiang, Saini Yi, Qin Liu, Jinqiang Zhang

## Abstract

Microglia has been reported to be able to regulate the proliferation, differentiation and survival of adult Neural stem/progenitor cells (NSPCs) by modulating the microenvironment, which results in different consequences of adult neurogenesis. However, whether the microglial activation is beneficial or harmful to NSPCs is still controversial because of the complexity and variability of microglial activation phenotypes. In this study, we detected the expression levels of M1 marker and M2 marker in IFN-γ- and IL-4-induced microglia at different time, respectively. The phenotypic markers of M1 and M2 microglia were stable for 24 h after removal of IFN-γ and IL-4 intervention, but exhibited different change patterns during the next 24 h. Then, the adult NSPCs were treated by the conditioned medium from IFN-γ- and IL-4-activated microglia. The conditioned medium from IFN-γ-activated microglia promoted apoptosis and astroglial differentiation of NSPCs, while suppressed proliferation and neuronal differentiation of NSPCs. However, the conditioned medium from IL-4-activated microglia exhibited opposite effects on these physiological processes. In addition, the direct treatment of IFN-γ or IL-4 alone did not significantly affect the proliferation, differentiation and survival of NSPCs. These results suggest that the secretome of pro-inflammatory (M1) and anti-inflammatory (M2) microglia differently regulated the proliferation, differentiation and survival of adult NSPCs. These findings will help further study the biological mechanism of microglia regulating neurogenesis, and provide a therapeutic strategy for neurological diseases by regulating microglial phenotypes to affect neurogenesis.

## 1. Introduction

Neural stem/progenitor cells (NSPCs) are multipotent cells that renew themselves and could differentiate into neurons and macro glia of the nervous system during embryonic development[1]. They are associated with learning, memory, and emotional behavior and play an important role in the neuropathological progression of mammals, including humans[2]. Therefore, regulating the function of adult NSPCs has great potential for the treatment of various neurodegenerative diseases. However, the neurobiological behaviors of NSPCs, such as proliferation or differentiation are affected by the environment where they reside. Thus, it is important to find effective ways to promote NSPCs’ proliferation, differentiation and survival.

Microglia are central nervous system (CNS)-resident immune cells and play a crucial role in the CNS by participating in local inflammatory reaction, regulating homeostasis of brain and microenvironment of neurogenesis, and modulating synapse formation[3]. It has been reported that microglia regulate proliferation, differentiation and survival of NSPCs by modulating the microenvironment where they reside[4]. However, there are considerable debates on whether microglial activation is favorable or unfavorable for NSPCs due to the complexity and variability of microglial activation phenotypes. Microglia are known to polarize into two reciprocate forms in response to external pro-inflammatory (M1) state and anti-inflammatory (M2) state[5]. M1 microglia disrupts the internal environment by release a variety of inflammatory factors, such as tumor necrosis factor alpha (TNF-α) and interferon γ (IFN-γ)[6]. In contrast, M2 microglia plays a neuroprotective role by secreting anti-inflammatory mediators and neurotrophic factors involved in homeostasis[7]. Modulation of microglial phenotype is an appealing neurotherapeutics.

Therefore, the objective of this study is to explore the activation phenotype of microglia was systematically investigated at different time after stimulation. Furthermore, we have investigated the effect of the secretome of different phenotype of microglia on the process of proliferation, neurogenesis and neural differentiation of NSPCs.

## 2. Materials and Methods

### 2.1. Microglia culture

Primary cultures of microglia cells were performed, as described previously[8]. In brief, the cerebral hemispheres were obtained from of neonatal (P0 - P3) C57BL/6J mice, minced tissue was enzymatically dissociated to a single cell suspension using 0.125% pancreatin, and the suspension was filtered with a 70 μm cell strainer. Filtrate after centrifugation prepared a cell suspension in DMEM/F12 containing 1% double antibodies and 10% FBS, incubated at 37 ° C, 5% CO_2_. Single microglia was collected on the 14th day by briefly shaking the mixed glial cultures. The procedures were approved by the Institutional Animal Care and Use Committee, University of Electronic Science and Technology of China.

### 2.2. Adult NSPCs culture

Adult NSPCs were obtained from the SVZ of 8-weeks-old C57BL/6J male mice accordingly to a protocol previously described[9]. Briefly, the entire SVZ region was dissected and the lateral wall of the lateral ventricles was carefully removed from the surrounding brain tissue into PBS. These tissues were chopped into 1 mm cubes and digested in 0.125% trypsin for 5 min, stopped from digestion with soybean trypsin inhibitor. The cells were re-suspended in DMEM/F12 complete medium containing 20 ng/mL recombinant murine FGF (PeproTech, 450-33), 20 ng/mL recombinant murine EGF (PeproTech, 315-09), 1% N2 (Gibico, 17502048) and 2% B-27 supplement (Gibico, 17504044). After culture for 7 days, the neurospheres were isolated by centrifugation (600 × g), enzymatically dissociated to a single cell suspension using 0.25% pancreatin, and plated at a density of 5 × 104 cells/cm^2^ in the proliferation medium. To permit serial cell passages, this pancreatin-dissociation process was repeated every 3 to 4 days.

### 2.3. Polarization of microglia

The microglia were plated at a density of 5 × 10 ^5^ cells /cm^2^ and treated with either IFN-γ (50 ng/mL, Sigma, SRP3211), IL-4 (20 ng/mL, Sigma, 11276905001) or PBS. Microglia were treated by IFN-γ or IL-4 for 24 h, 48 h and 72 h, and the area of each microglia was measured by image J. Activated microglia were stained by immunofluorescence, and the genes and protein expression levels of M2 markers (Arg-1, YM-1, CD206 and IL-10) and M1 marker (INF-α, iNOS, IL-6 and CCR2) were detected at 24 h, 48 h and 72 h after treatment.

### 2.4. NSPCs conditional culture and co-culture with microglia

The microglia were plated at a density of 5 × 10^5^ cells/cm^2^ and treated with either IFN-γ or IL-4 for 24h, microglia cells were washed by PBS twice then added DMEM/F12 + GlutaMax culture medium for 24 h. The microglial medium supernatant was collected and made conditioned medium for differentiation and proliferation of NSPCs. NSPCs cultured in different conditioned medium from PBS-, IL-4- or IFN-γ-treated microglia and control medium added PBS, IFN-γ and IL-4. The proliferation, differentiation and survival of NSPCs were detected by immunofluorescence staining. The neurosphere co-cultured with microglia for 24 h by Transwell as previous described.

### 2.5. Immunofluorescence

The cells (including microglia, neurosphere, proliferative NSPCs and differential NSPCs) were plated at a density of 1 × 10 ^5^ cells/cm^2^ and fixed with 4% paraformaldehyde (PFA; pH 7.2) for 30 min. The cells were permeabilized with 0.5% Triton X-100 in PBS for 15 min. Then the cells were blocked in 10% donkey serum for 2 h and incubated with primary antibodies (goat anti-Iba1, 1:400, Abcam; mouse anti-CD68, 1:400, Cell Signaling Technology; rabbit anti-Arg-1, 1:400, Abcam; rabbit anti-iNOS, 1:50, Abcam; goat anti-DCX, 1:400, Santa; mouse anti-BrdU, 1:400, Cell Signaling Technology; mouse anti-GFAP, 1:400, Cell Signaling Technology; anti-CC3, 1:150, Cell Signaling Technology) overnight at 4°C, and further incubated with fluorescent-dye-conjugated secondary antibodies (DyLight 549-conjugate donkey anti-goat, 1:300, Jackson ImmunoResearch; DyLight 488-conjugate donkey anti-mouse, 1:300, Jackson ImmunoResearch) for 2 h at room temperature. Subsequently, the cells were incubated with DAPI (1:10000, Roche) for 5 min and imaged using a fluorescent microscope (Olympus BX51). The Iba1+ area and CD68^+^ area was quantized by Image J software (version 1.45 J; National Institutes of Health, Bethesda, MD, USA).

### 2.6. Real-time polymerase chain reaction (RT-PCR)

Cell plates of each group were selected and washed with PBS. Total ribonucleic acid (RNA) was isolated by using the Trizol reagent, according to the standard program. RT-qPCR was performed using a First Strand cDNA Synthesis Kit (TaKaRa), according to the manufacturer’s instructions. The threshold amplification cycle number (Ct) was determined for each reaction by Quantitative RT-PCR (Bio-Rad CFX 96) in the linear phase of the amplification plot. Each sample was tested in triplicate, and an analysis of gene expression was performed using the -ΔΔCt method. The values were normalized against the housekeeping genes β-actin. The primer sequences of each gene are described in Table 1.

**Table 1.**
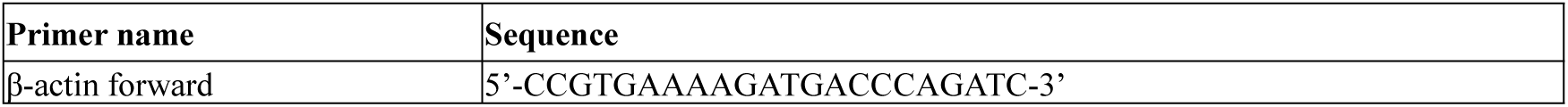
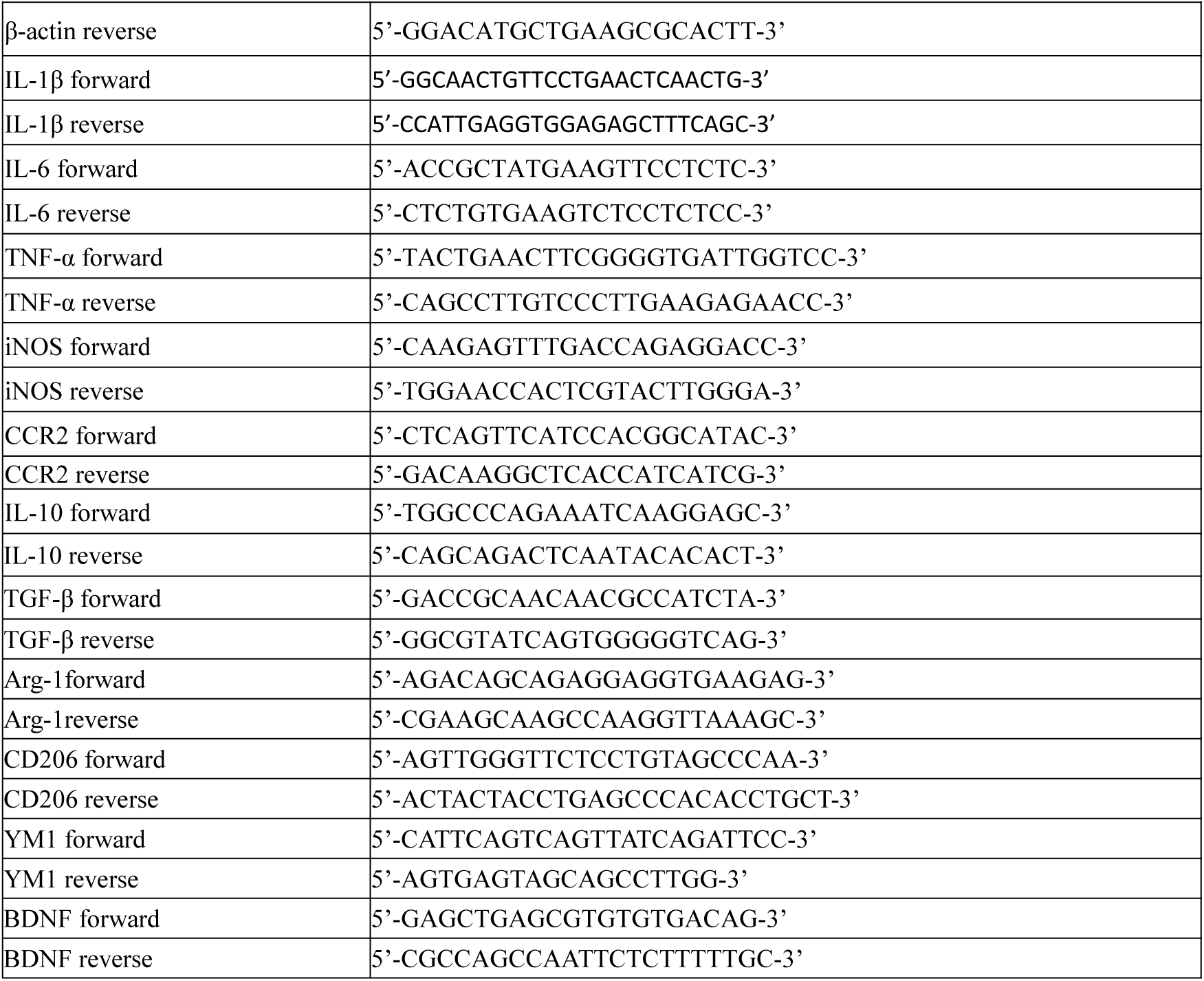
Primers used in quantitative RT-PCR.

### 2.7. ELISA

Enriched microglia were cultured in 6-well culture plate at 5 × 10 ^5^ cells/cm^2^, and treated with either IFN-γ, IL-4 or PBS. 24 h or 48h after continuous stimulation or cessation of stimulation, the medium was collected, and the enriched microglia were fractured by a cell lysis buffer (Solarbio, China), and the lysates were centrifuged at 1,000 × g for 30 min. The supernatants were used to examine the cytokine concentration. The concentrations of Arg-1, iNOS, TNF-α and TGF-β were quantified using ELISA kits (QuantiCyto, China) according to the manufacturer’s protocol. The detection limit for them was 8 pg / mL.

### 2.8. Statistical analysis

All of the data were expressed as means ± SEMs. The statistical analyses were performed using Graghpad prism (version 5; GraphPad). Individual comparisons were assessed using Student’s two-tailed t test, and multiple comparisons were performed with two way or one-way analysis of variance and Tukey’s post hoctests. Statistically significant differences were defined as *P* < 0.05.

## 3. Result

### 3.1. M1-like and M2-like microglia were treated by IFN-γ or IL-4

First, we assessed the shape and area of microglia with different phenotypes. The untreated microglia were fusiform, while the microglia induced by IFN-γ exhibited oval shape and thorny convexity, and the microglia activated by IL-4 were multiramose (Fig. 1A). In addition, the results show that the area of microglia stimulated with IL-4- or IFN-γ were larger than the area of untreated microglia after treated by IFN-γ or IL-4 for 24 h, 48 h and 72 h (Fig 1B).

**Figure 1:**
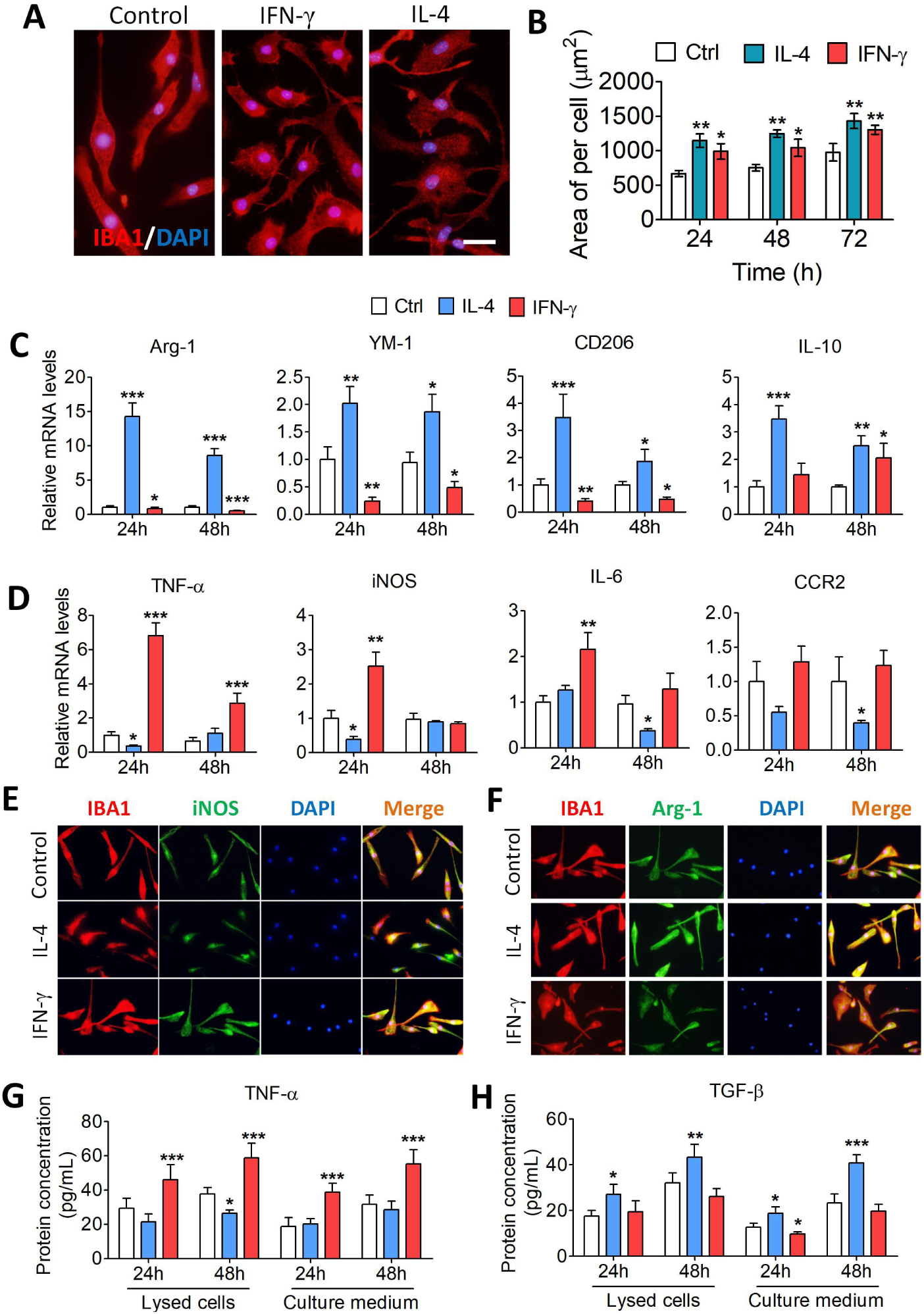
Characterization of microglia under IFN-γ or IL-4 intervention conditions. (A) The morphological micrograph for microglia when exposed in PBS, IFN-γ or IL-4 for 24 h. The microglia were stained IBA1 (red) using immunocytochemical staining and the nucleus is labeled by DAPI (blue). Scale bar is 10 μm. (B) The bar graph shows the area of each microglia. (C and D), The M2 and M1 markers expression of the microglia exposed in PBS, IFN-γ or IL-4 for 24 h and 48 h. (E and F) Micrograph show the expression of M1 markers (iNOS, green) and M2 markers (Arg-1, green) when exposed in PBS, IFN-γ or IL-4 for 24 h. The microglia were stained IBA1 (red) and the nucleus is labeled by DAPI (blue). Scale bar is 10 μm. (G and H) The bar graph shows the concentration of TNF-α and TGF-β both inside and outside the microglia exposed in PBS, IFN-γ or IL-4 for 24 h, 48 h. Data are showed Mean ± SEM, n = 4-6, \**P* < 0.05, \*\**P* < 0.01, \*\*\**P* < 0.005 vs control group.

The RT-PCR results confirmed that IL-4 and IFN-γ had opposite effects on the activation of microglia, and the responses of M1 markers and M2 markers to either stimulus were also different. Specifically, after treatment with IL-4 for 24 h and 72 h markedly increased the expression of M2 markers (Arg-1, YM-1, CD206 and IL-10) and decreased expression of M1 markers (INF-α, iNOS, IL-6 and CCR2) in microglia (Fig. 1C). In contrast, we found that the expression of M2 markers (Arg-1, YM-1 and CD206) was decreased while the expression of M1 markers (INF-α, iNOS and IL-6) was increased after treatment IFN-γ for 24 h and 48 h (Fig. 1D).

The result of protein expression showed that the IL-4 treatment decreased the extracellular expression of iNOS in microglia, but increased the expression of Arg-1. Meanwhile, an opposite trend was observed in IFN-γ-activated microglia (Figs. 1E and 1F). The result of ELISA indicated that the protein concentrations of intracellular and extracellular TNF-α in IFN-γ-activated microglia were significantly higher than those in untreated microglia either at 24 h or 48 h after stimulation. While the protein concentrations of TGF-β in IL-4-treated microglia were significantly higher than those in untreated microglia in both 24 h and 48 h after stimulation (Figs. 1G and 1H). These results demonstrated that the M1 or M2 phenotype of microglia induced by IFN-γ or IL-4 respectively was temporarily stable between 24 h and 48 h after stimulation.

### 3.2. Phenotypic maintenance of M1 and M2 microglia after removal of IFN-γ or IL-4 intervention

Then we assessed the gene expression of M2 markers and M1 marker in activated microglia at 24 h and 48 h after the intervening factors were removed (Fig. 2A). The data show that IL-4-treated microglia had higher M2 markers expression (Arg-1, and TGF-β) at 24 h but no significant change at 48 h after removal of intervention when compared to untreated microglia. Increases of M1 marker expression (TNF-α, iNOS and IL-6) and M2 marker expression (Arg-1), relative to the untreated microglia, were recorded in INF-γ-treated microglia at 24 h and 48 h after removal of intervention, respectively (Figs. 2B and 2C). The intracellular protein concentrations of iNOS, Arg-1 and TGF-β in IFN-γ- and IL-4-treated microglia were assessed at 0 h, 24 h and 48 h after removal of IFN-γ or IL-4 intervention using ELISA. The results were consistent with gene expression level (Figs. 2D).

**Figure 2:**
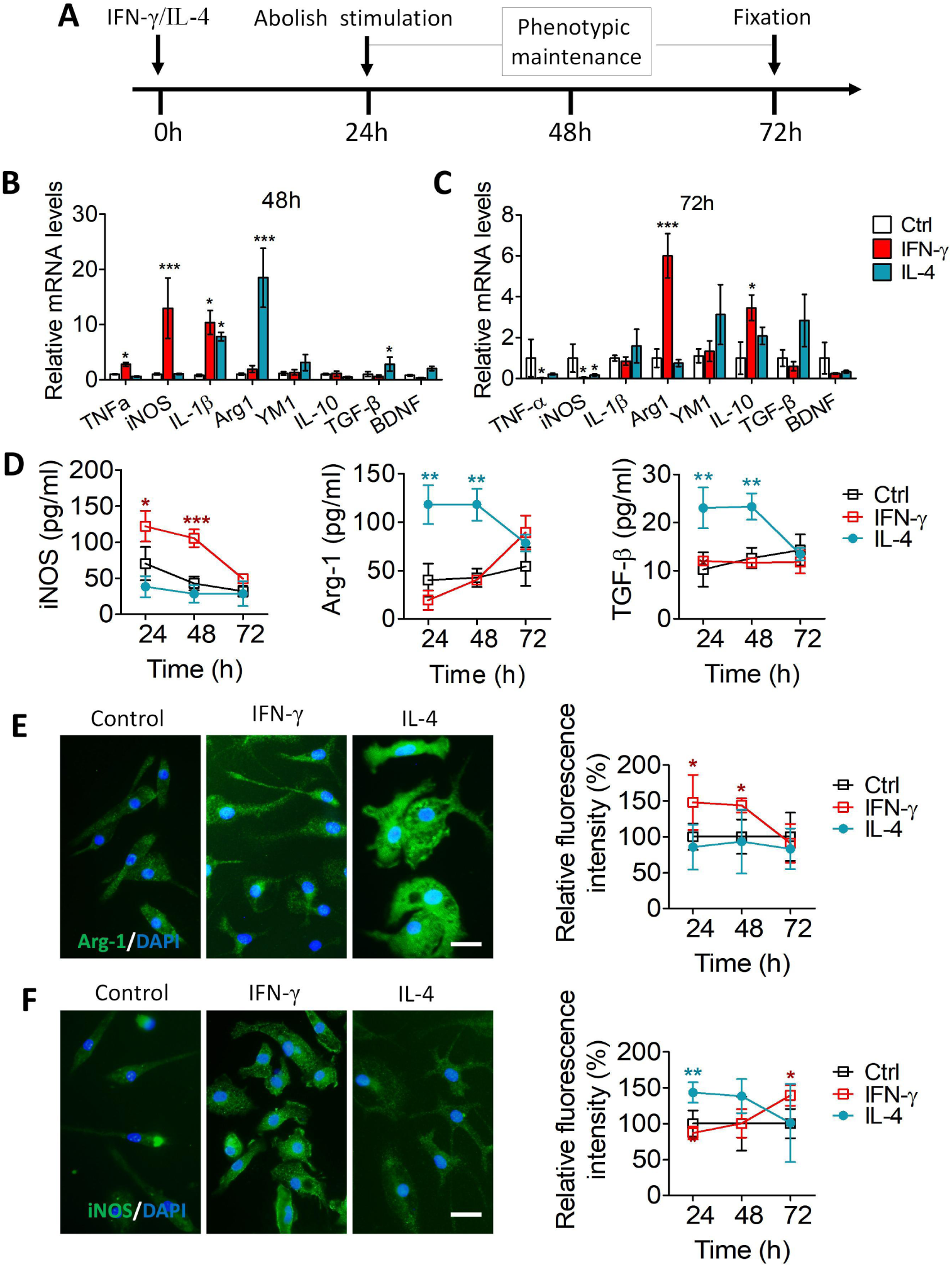
Phenotype maintenance of M1 and M2 microglia after removing IFN-γ or IL-4 intervention. (A) The scheme for explore the time of microglia maintain phenotypic duration after remove IFN-γ or IL-4 intervention. After microglia activated by IFN-γ and IL-4 for 24 h, IFN-γ and IL-4 were replaced with serum-free culture medium, and microglia were cultured for another 24 h and 48 h. (B and C) The M2 and M2 markers expression of the active microglia (treated by IFN-γ or IL-4) when removed intervention for 24 h and 48 h. (D) The line graph illustrate the tendency of expression of M1 markers (iNOS) and M2 markers (Arg-1 and TGF-β) after remove intervention for 24 h and 48 h. (E and F) The immunofluorescent staining of Arg-1 and iNOS in microglia treated by PBS, IFN-γ or IL-4 when remove intervention for 24 h. And the line graph shows the trend of expression of Arg-1 and iNOS when remove intervention for 24 h and 48 h. Data are showed Mean ± SEM, n = 4-6, \**P* < 0.05, \*\**P* < 0.01, \*\*\**P* < 0.005 vs control group.

In addition, the results indicated that in comparison with the untreated microglia the expression of Arg-1 and iNOS increased in IL-4 and IFN-γ treated microglia respectively at 24 h after removal of intervention, but both of them fell back to normal after another 24 h (Fig. 2E and 2F). It is worth mentioning that Arg-1 was significantly upregulated in IFN-γ-activated microglia at 48 h after removal of intervention.

### 3.3. Effects of the secretome from M1 and M2 microglia on proliferation of adult NSPCs

The NSPCs were isolated from brain of adult mice (Fig. 3A), and were treated with PBS, IL-4 or IFN-γ, respectively, or co-cultured with PBS-, IFN-γ- or IL-4-treated microglia for 24 h (Fig. 3B). There was no significant difference in the neurosphere size when adult NSPCs were exposed to PBS, IL-4 and IFN-γ alone. However, compared to those co-cultured with PBS-treated microglia, the neurosphere size increased when adult NSPCs co-cultured with IL-4-induced microglia, and decreased when co-cultured with IFN-γ-induced microglia (Figs. 3C and 3D).

**Figure3:**
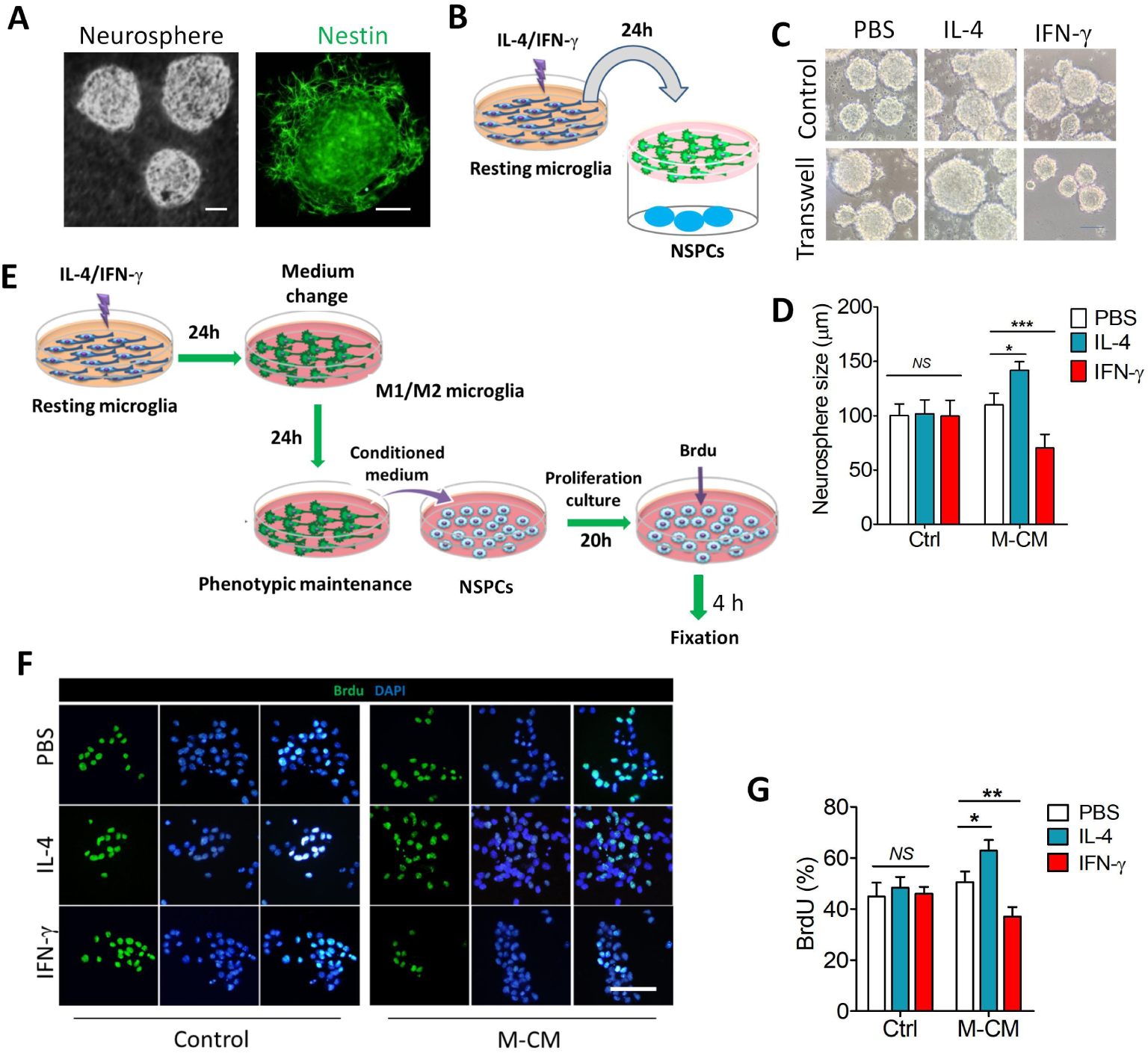
Effects of the secretome from M1 and M2 microglia on proliferation of adult NSPCs. (A) The micrograph shows the suspended growth of neurosphere for four days. The immunostaining depicted isolated cells expressing Nestin (green). Scale bar is 10 μm. (B) The NSPCs were co-cultured with PBS-, IFN-γ- or IL-4-treated microglia for 24 h. (C and D) The changes of size of neurosphere (E) The program flowchart for studying the effects of the secretome from M1 and M2 microglia on proliferation of adult NSPCs. (F and G) The percentage of BrdU^+^ cells from NSPCs co-cultured with microglia induced by PBS, IL-4 or IFN-γ for 24 h. Data are showed Mean ± SEM, n = 4-6, *P < 0.05, **P < 0.01, ***P < 0.005 vs control group.

To further confirm the effects of IL-4 or IFN-γ activated microglia on the proliferation of adult NSPCs, the NSPCs were separately cultured in the conditioned medium (CM) collected from PBS, IFN-γ or IL-4 treated microglia. The proliferation of NSPCs was assessed by labeled with BrdU (Fig. 3E). There was no significant change in the percentage of BrdU+ cells when adult NSPCs were exposed to PBS, IL-4 or IFN-γ alone. However, the percentage of BrdU+ cells increased when the adult NSPCs were cultured in the CM from IL-4-treated microglia, but decreased when cultured in the CM from INF-γ-treated microglia (Fig.4F and 4G). These results suggest that although INF-γ and IL-4 exert no influence on NSPCs proliferation, the factors secreted by the INF-γ-treated microglia can negatively affect NSPCs proliferation, while the factors secreted by the IL-4-treated microglia are beneficial to NSPCs proliferation.

**Figure 4:**
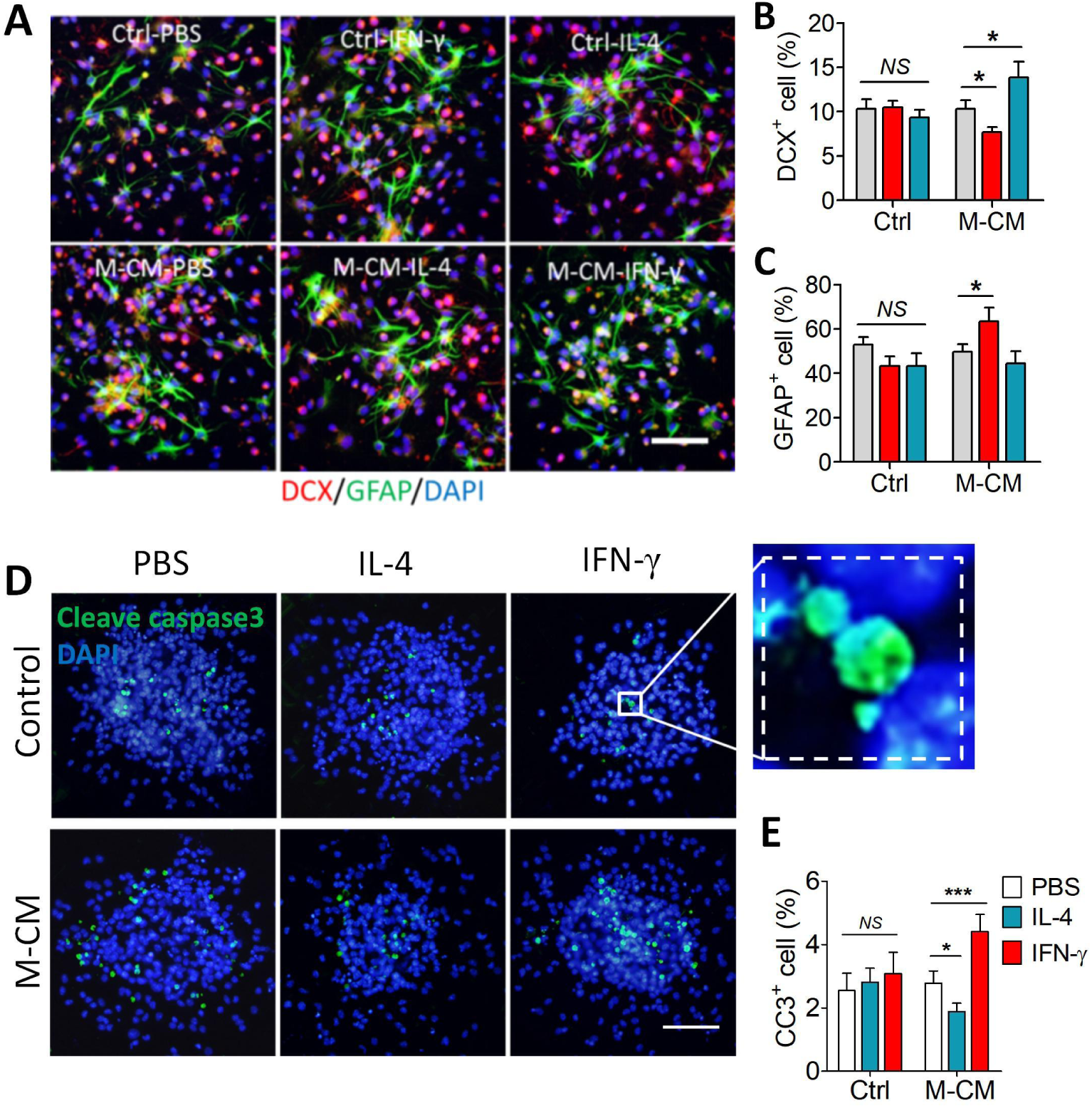
Effects of the secretome from M1 and M2 microglia on differentiation and survival of adult NSPCs. (A) The morphological micrograph for neurosphere when differentiate for 3 days in PBS, IFN-γ or IL-4 for 24 h, or in CM from microglia induced by PBS, IL-4 or IFN-γ for 24 h. Scale bar is 10 μm. (B) The bar graph shows the percentage of DCX^+^ cells from NSPCs differentiate for 3 days in CM microglia induced by PBS, IL-4 or IFN-γ for 24 h. (C) The bar graph shows the percentage of GFAP^+^ cells from NSPCs differentiate for 3 days in CM from microglia induced by PBS, IL-4 or IFN-γ for 24 h. (D) The morphological micrograph for NSPCs when exposed in PBS, IFN-γ or IL-4 for 24 h, or co-cultured with microglia induced by PBS, IL-4 or IFN-γ for 24 h. The apoptotic cells were stained Cleave caspase3 (green) using immunocytochemical staining and the nucleus is labeled by DAPI (blue). Scale bar is 10 μm. (E) The bar graph shows the percentage of CC3^+^ cells from NSPCs co-cultured with microglia induced by PBS, IL-4 or IFN-γ for 24 h. Data are showed Mean ± SEM, n = 4-6, \**P* < 0.05, \*\**P* < 0.01, \*\*\**P* < 0.005 vs control group.

### 3.4. Effects of the secretome from M1 and M2 microglia on differentiation and survival of adult NSPCs

To determine the role of activated microglia in functional neurogenesis, we analyzed the percentage of cells committed to neuron or astrocyte which differentiate in CM from IFN-γ-induced M1 microglia and IL-4-induced M2 microglia for 3 days. The results indicate that exposure to PBS, IL-4 or IFN-γ alone had no significant effect on the percentage of DCX^+^ or GFAP^+^ cells. However, the percentage of DCX^+^ cells decreased in the CM from the IFN-γ-treated microglia, but increased in the CM from the IL-4-treated microglia (Fig. 4A and 4B). In contrast, the percentage of GFAP^+^ cells increased in the CM from the IFN-γ-treated microglia, while decreased in the CM from the IL-4-treated microglia (Fig. 4A and 4C). These results suggest that factors secreted from the IFN-γ-treated microglia benefit astrocytes differentiation compared to the untreated microglia, while the factors secreted from the IL-4-treated microglia can promote neurons differentiation.

Moreover, the results reveal that PBS, IL-4 and IFN-γ exerted no significant influence on the percentage of CC3^+^ cells. However, the percentage of CC3^+^ cells in the CM from IL-4 or INF-γ treated microglia decreased or increased respectively compared to the untreated microglia (Fig. 4D and 4E). These results suggest that factors secreted from the INF-γ or IL-4 treated microglia can negatively or positively influence NSPCs survival, despite that INF-γ or IL-4 itself has no significant effect on NSPCs survival.

## 4. Discussion

Microglia are myeloid cells that regulate proliferation, differentiation and survival of NSPCs by modulating the microenvironment where they are located[10]. However, there has been considerable debate as to whether microglial activation is favorable or unfavorable for NSPCs. The complexity and variability of microglial activation phenotypes are responsible for this confusion[11]. In this study, we determined the activation phenotype of microglia at different times after stimulation. Moreover, we have further demonstrated that proliferation, neural differentiation and survival of adult NSPCs were suppressed by the secretome of M1 microglia, but were promoted by the secretome of M2 microglia. To our knowledge, this is the first time to identify and provide a clear result for the influence of different phenotypic microglia on the proliferation, neural differentiation and survival of adult NSPCs.

Microglia showed strictly stimuli dependent and related to the pathological conditions[12]. In this study, microglia stimulated continuously by IL-4 for 24 h and 48 h showed multiramose, higher expression level of M2 markers and lower expression level of M1 markers compared to the untreated microglia. In contrast, those treated continuously with INF-γ for 24 h and 48 h showed oval shape and thorny convexity, significant higher expression level of M1 markers and low expression level of M2 markers. These results suggest that microglia can polarize to M2 microglia or M1 microglia by IL-4 and INF-γ, respectively. And their activation can persist for a long time as a double-edged sword. However, whether microglia treated by IL-4 or IFN-γ can maintain M1 or M2 phenotype after removal of the stimulus intervention is less well-defined. Previous research showed that 24 h-LPS-treatment induced M1 phenotypic microglia, whereas 72 h-LPS-treatment induced M2 phenotypic microglia in vitro, which indicated that the activation state of microglia was significantly dependent on the stimulation duration[13]. In the present study, expression of M2 markers (Arg-1, and TGF-β) in IL-4-treated microglia increased during the first 24 h after stimulus removal, but fell back to normal within the next 24 h. Nevertheless, compared to the untreated microglia, higher expression of M1 marker (TNF-α, iNOS and IL-6) and M2 marker (Arg-1) was observed in INF-γ-treated microglia at 24 h and 48 h after removing the stimulation, respectively. These data suggest microglia maintain a relatively stable phenotype (M1 or M2) within 24 hours after removal of stimulus.

Neural stem/progenitor cells (NSPCs) undergo a series of developmental processes before giving rise to newborn neurons, astrocytes and oligodendrocytes in adult neurogenesis[14]. It has been proven that microglia regulate the proliferation, differentiation and survival of adult NSPCs in different ways, which results in the varying consequences of adult neurogenesis[15]. However, the effects of M1 and M2 microglia on neural stem cells are controversial, due to the complexity of the microglia phenotype, especially after removal of stimulus intervention. In this study, the conditioned medium from IFN-γ-activated microglia was found to suppress the proliferation, differentiation and survival of NSPCs. While the conditioned medium from IL-4-activated microglia showed an opposite effect on NSPC proliferation and differentiation. Moreover, the direct treatment of IFN-γ or IL-4 alone did not significantly affect the proliferation, differentiation and survival of NSPCs. So, at the first 24 hours, M1 microglia induced by IFN-γ can suppress the proliferation, differentiation and survival of NSPCs through releasing M1 markers such as NO, IL-1β, TNF-α, etc. Then at 48 h after the cessation of stimulation, M2 microglia become immune-inactivated, while M1 microglia may transform into M2 microglia. And M2 microglia can positively affect the aforementioned processes of NSPCs by secreting M2 markers including BDNF, TGF-β, IGF-1, etc. In addition, recent work has indicated that the astrocytes-derived factors showed suppression of NSPC proliferation[9]. In this study, the stimulation secreted from the IFN-γ-treated microglia benefit astrocytes differentiation. This result may also partially explain the negative effect of M1 microglia on the proliferation and differentiation of NSPC.

In conclusion, our results suggest that how long M1 and M2 microglia maintain their phenotype after stimulus removal. And we also demonstrated a bidirectional regulation of neuronal cells by secretions from M1 and M2 microglia. These findings will help further study the biological mechanism of microglia regulating neurogenesis, and provide a therapeutic strategy for neurological diseases by regulating microglial phenotypes to affect neurogenesis.

## Acknowledgment

This work was supported by the National Natural Science Foundation of China (82060726).

## Notes

### Competing Interest Statement

The authors have declared no competing interest.

